# Modulating Cas13a *trans*-cleavage by double strand RNA: Application to the development of an autocatalytic sensor

**DOI:** 10.1101/2024.03.05.583472

**Authors:** Fei Deng, Sneha Gulati, Rui Sang, Yi Li, Ewa M. Goldys

## Abstract

Cas13a-based diagnostic systems have been widely utilized for the detection of RNA targets. However, without preamplification such systems have sensitivity in the picomolar range only. Here, we found that double strand RNA (dsRNA) over 20nt is able to effectively activate the *trans*-cleavage activity of Cas13a RNP, while the cleavage rates of dsRNA by activated Cas13a RNP are very low. In addition, specially designed small circular RNA constructs (Cir-mediators comprising a 20nt dsRNA trigger with a 5nt ssRNA linker) have limited ability to activate Cas13a RNP, but this activation is restored once the circular structures are cleaved and become linear. Based on this new method to control *trans*-cleavage activity of Cas13 RNP, we developed a Cas13a autocatalytic biosensing system assisted by Cir-mediators, which allows one target RNA to activate numerous Cas13a RNPs. With this approach we show ultrasensitive detection of 1aM of synthetic RNA targets without preamplification within 15min. The sensor was successfully applied to monitor miRNA-21 concentration in clinical plasma samples in colorectal cancer. This investigation yields novel insights into the properties of Cas13a RNPs, and the Cir-mediator-based autosensor introduces a novel method for detecting RNA targets with exceptional sensitivity.

**Highlights:**

1. dsRNA is able to trigger activation of Cas13a RNP.
2. Activated Cas13a RNP do not cleave dsRNA.
3. Cir-mediator induces low levels of Cas13a RNP activation.
4. Cir-mediator based Cas13a auto-catalysis biosensor can detect 1aM RNA targets.

## Introduction

Cas13a is a programmable RNA-guided ribonuclease^1, 2^. When bound to a target RNA, Cas13a displays both sequence-specific RNA target cleavage (*cis*-cleavage) and collateral cleavage (*trans*-cleavage) characteristics^1, 2^. The *trans*-cleavage is realised by the HEPN catalytic site revealed upon Cas13a activation which is capable of non-specific cleavage of single-strand RNA (ssRNA)^2^. This can be exploited to create sequence-specific RNA biosensors, however basic Cas13a biosensing systems which recognise the presence of RNA targets by cleaving ssRNA fluorescent quenched reporters show limited sensitivity in the pM range^3, 4^. Thus nucleic acid preamplification techniques have been required to realize ultrasensitive (aM level) detection of RNA targets in Cas13a systems, such as the recombinase polymerase amplification (RPA) in the SHERLOCK,^3, 5^ and LAMP in the DISCoVER^6^ systems. These nucleic acid amplification methods require specialised laboratories/equipment, and they also suffer from false positive readings^7^ and amplicon contamination^8^. The requirement for amplification currently hinders the creation of convenient, low-cost assays with the requisite specificity and sensitivity for nucleic acid analysis in clinical settings. Therefore, there is a demand for sequence-specific detection methods of RNA targets that do not rely on amplification.

Up until now, two primary types of approaches for detecting RNA targets without amplification have been reported. One type of approach is to integrate diverse signal amplification techniques, such as polydisperse droplet digital assay^9^, mobile phone microscopy^10^, nanoenzyme^11^, or field-effect transistors^12^ with CRISPR/Cas systems. These methods either require specialised instrumentation or are difficult to realize a point-of-care setting. Another type of approach is to employ the Cas tandem system, where the first Cas protein performs target recognition, and a second Cas protein realises cascade signal amplification; examples include Cas13-Csm6 ^3, 13^, Cas13-Cas14^14^, and Cas13-Cas12 systems^15, 16^. In comparison with a basic Cas13a system, where one target activates one Cas13a RNP, the Cas tandem systems are able to achieve higher sensitivity. In these systems one target activates 10^2^-10^3^ Cas RNPs^13-16^, leading to increased signal amplification. An autocatalytic reaction is expected to further increase biosensing performance, because one target will, in principle, be able to activate unlimited Cas RNPs^17, 18^. To date, a Cas13 autocatalytic reaction system has not been reported.

In this study, we investigated fundamental properties of Cas13a nuclease, including the ability of ssRNA, dsRNA and circular RNA to activate the Cas13a RNP as well as the *trans*-cleavage activity of Cas13a on ssRNA, dsRNA, and circular RNA substrates. We found that dsRNA is able to activate a Cas13a RNP, while such activated Cas13a RNP was not able to cleave dsRNA. In addition, circular RNA could activate Cas13a only to a very limited degree, while after linearization the activation property has been recovered. Based on these findings, we developed a new type of autocatalytic biosensor (termed “autosensor”) based on Cas13a. Its key component is a molecular construct called “Cir-mediator” which consists of a dsRNA region acting as a replicate activation trigger for Cas13a, and a short ssRNA linker which controls the activation property of dsRNA. The Cir-mediator facilitates the autocatalysis reaction in the following way. In the presence of a target RNA molecule, one Cas13a RNP is activated, which continues to cleave the ssRNA region of the supplied Cir-mediators to release the dsRNA triggers which act as surrogates of the original detection targets. These triggers then activate new Cas13a RNPs. In this way, one target can theoretically activate unlimited number of Cas RNPs in the reaction system. We show here that the optimized Cas13a autosensor is able to detect 1aM miRNA targets from clinical samples within 15min. Our autosensor provides a novel platform for POC testing of RNA targets without preamplification.

## Results

### 1. Double strand RNA is able to trigger activation of Cas13a RNP

To date, Cas13a has been reported to be effectively activated by single strand RNA (ssRNA) (Fig. S1)^1^. To explore alternative triggers of Cas13a, we investigated the previously unexplored types of single strand nucleic acids targets such as single strand DNA (ssDNA) and phosphorothioate-modified single strand RNA (ps-ssRNA). As shown in Fig.1B & Fig.S2, ssDNA was unable to activate Cas13a, while a slight signal increase was observed on ps-ssRNA (7.5%). A range of double strand nucleic acids were also investigated (Fig.1C & Fig.S2), including double strand RNA (ssRNA/cRNA), RNA/DNA hybrid trigger (ssRNA/cDNA), double strand DNA (ssDNA/cDNA), and ps-ssRNA/DNA hybrid trigger. In comparison with ssRNA trigger, the dsRNA target shows a significant level of Cas13a activation (53%), while the RNA/DNA hybrid target shows limited activation (14.3%), and no activation was observed for double strand DNA and ps-ssRNA/DNA hybrid targets. Therefore, our explorations reveal that while ps-ssRNA and RNA/DNA hybrid target produce limited activation, double strand RNA (dsRNA) unexpectedly emerged as a highly effective activator of Cas13a.

To investigate the Cas13a RNP activation mechanism by dsRNA, a FRET approach was applied^19^. To this aim, as shown in Fig. 1D, the two strands in dsRNA were labelled on one end with a fluorophore and a quencher (t-RNA with FAM, c-RNA with BHQ1); in such conditions limited background fluorescence signal was observed. The interaction of this dsRNA with Cas13a RNP, is expected to involve binding of the t-RNA to gRNA of Cas13a RNP, and release of the cRNA, which should increase the FRET signal. This was verified in Fig. 1E, where a low fluorescence signal of dsRNA was observed, while the signal for ssRNA-FAM was significantly higher. The fluorescence signal of Cas13a RNP upon interaction with dsRNA was comparable to that of ssRNA-FAM, confirming the binding of t-RNA on the gRNA of Cas13a.

**Figure 1.**
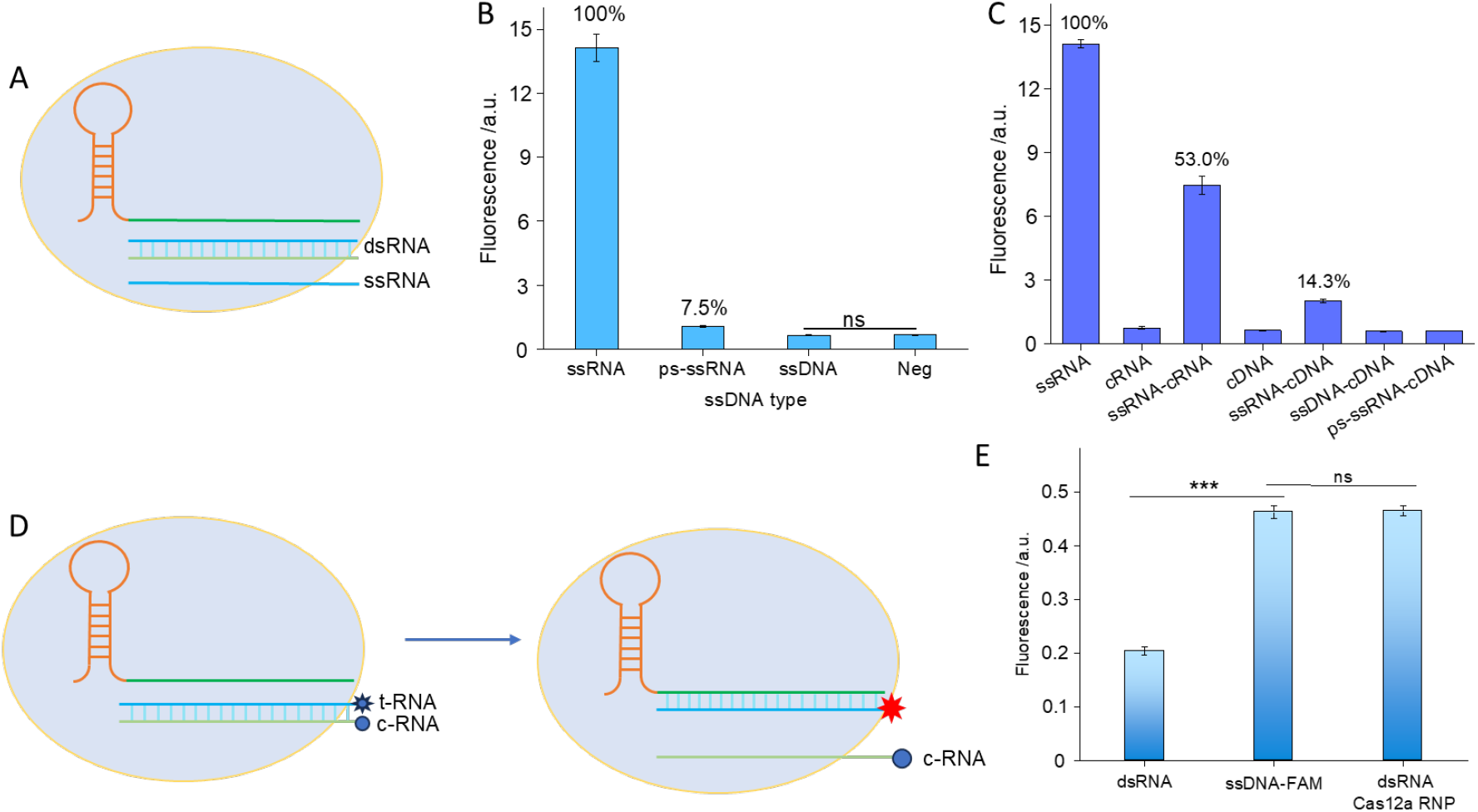
Investigation of different types of activation triggers for Cas13a RNP. (A) Schematic of a single strand or double strand trigger of Cas13a RNP; (B) Investigation of different types of single strand trigger, including ssRNA, ps-ssRNA and ssDNA (n=3); (C) Investigation of different types of double strand trigger, including dsRNA, RNA/DNA, dsDNA and psRNA/DNA (n=3); (D) Schematic of the FRET approach for the investigation of the dsRNA Cas13a activation mechanism; (E) FRET measurements shed light on the dsRNA Cas13a activation mechanism confirming the binding of t-RNA on the gRNA of Cas13a (n=3). (* P<0.05, ** P< 0.005, *** P< 0.001)

### 2. Activated Cas13a RNP lacks the ability to perform *trans*-cleavage of dsRNA

Published reports indicate that Cas13a RNP is capable of *trans*-cleavage of ssRNA (Fig. S3) ^2^. However, there has been relatively little investigation into the *trans*-cleavage activity of Cas13a on dsRNA substrates explored in this section (Fig. 2A). As shown in Fig. 2B, agarose gel electrophoresis shows that dsRNA is not cleaved by activated Cas13a. To further confirm this finding, a double strand fluorescent reporter was applied in a CRISPR/Cas13a biosensing system (Fig. 2C), and no significant fluorescence increase was seen, demonstrating that dsRNA cannot be cleaved by activated Cas13a.

**Figure 2.**
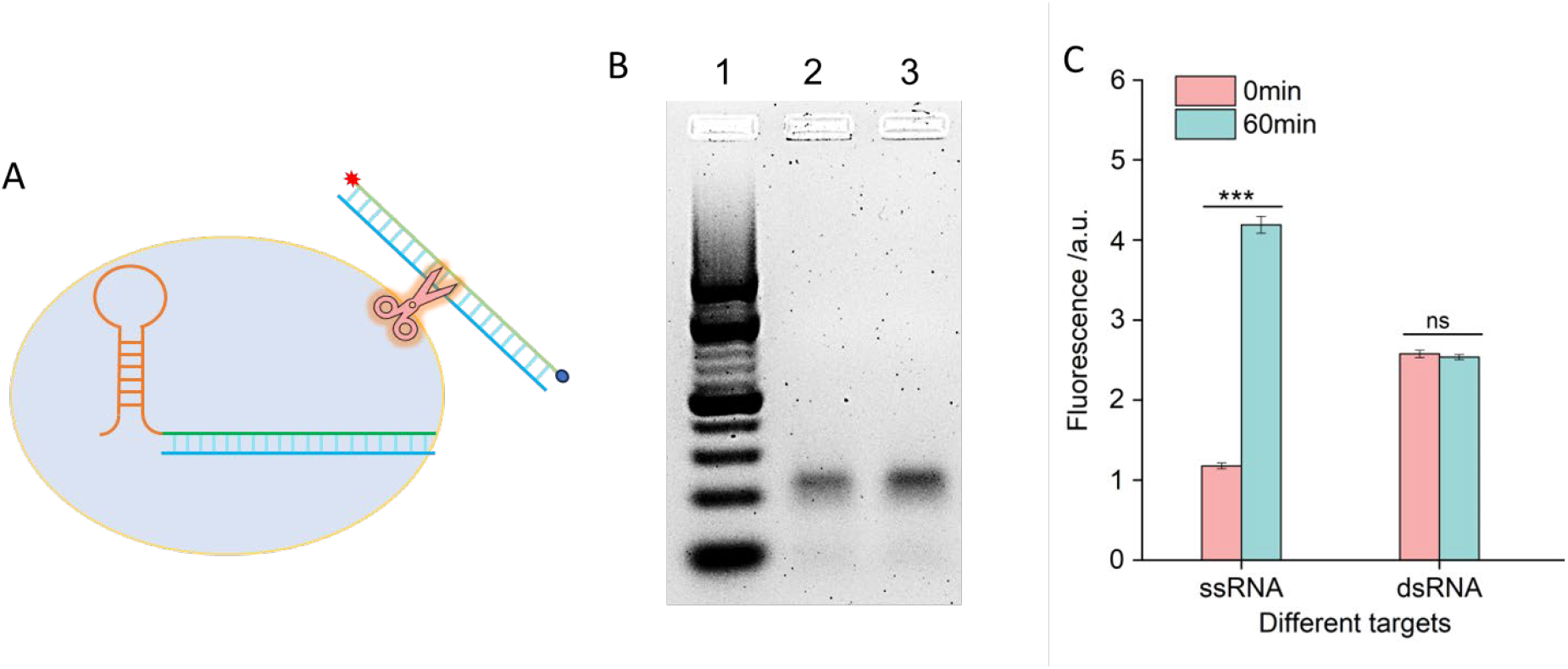
Investigation of the *trans*-cleavage ability of Cas13a RNP on dsRNA. (A) Schematic of the *trans*-cleavage activity of Cas13a RNP on dsRNA; (B) Investigation of the *trans*-cleavage activity of Cas13a on dsRNA substrates using gel electrophoresis assay. 1). 10 bp ladder; 2). dsRNA; 3) CRISPR/Cas13a treated dsRNA; (C) Investigation of the *trans*-cleavage activity of Cas13a on ssRNA and dsRNA substrates using fluorescent reporters (n=3). (* P<0.05, ** P< 0.005, *** P< 0.001)

### 3. Synthesis and characterization of Cir-mediator

Since dsRNA is an effective activation trigger of Cas13a RNP, but not a *trans*-cleavage substrate of activated Cas13a RNP, we constructed a Cir-mediator structure comprising a double strand RNA section and a short fragment of ssRNA (Fig. 3). The Cir-mediator was synthesised by using circular ssRNA with a slightly shorter cRNA. The circular ssRNA (Cir-ssRNA) was synthesized by using a click chemistry method based on our previous work^18^, and it was characterised by agarose gel electrophoresis (Fig. 3A & S4). As shown in Fig. 3A, Cir-ssRNA moves more slowly than linear ssRNA, confirming a difference in molecular conformation. In addition, reduced activation of Cas13a by Cir-ssRNA compared with linear ssRNA was observed (Fig. 3B), consistent with circular conformation of Cir-ssRNA. After mixing Cir-ssRNA with its cRNA, Cir-dsRNA was formed. Lower fluorescence signals for Cir-dsRNA were observed in Fig. 3C compared to the corresponding linearised structures, indicating that circular targets were locked by the linker with limited ability to trigger activation of Cas13a, and shorter linker length led to lower background signals.

**Figure 3.**
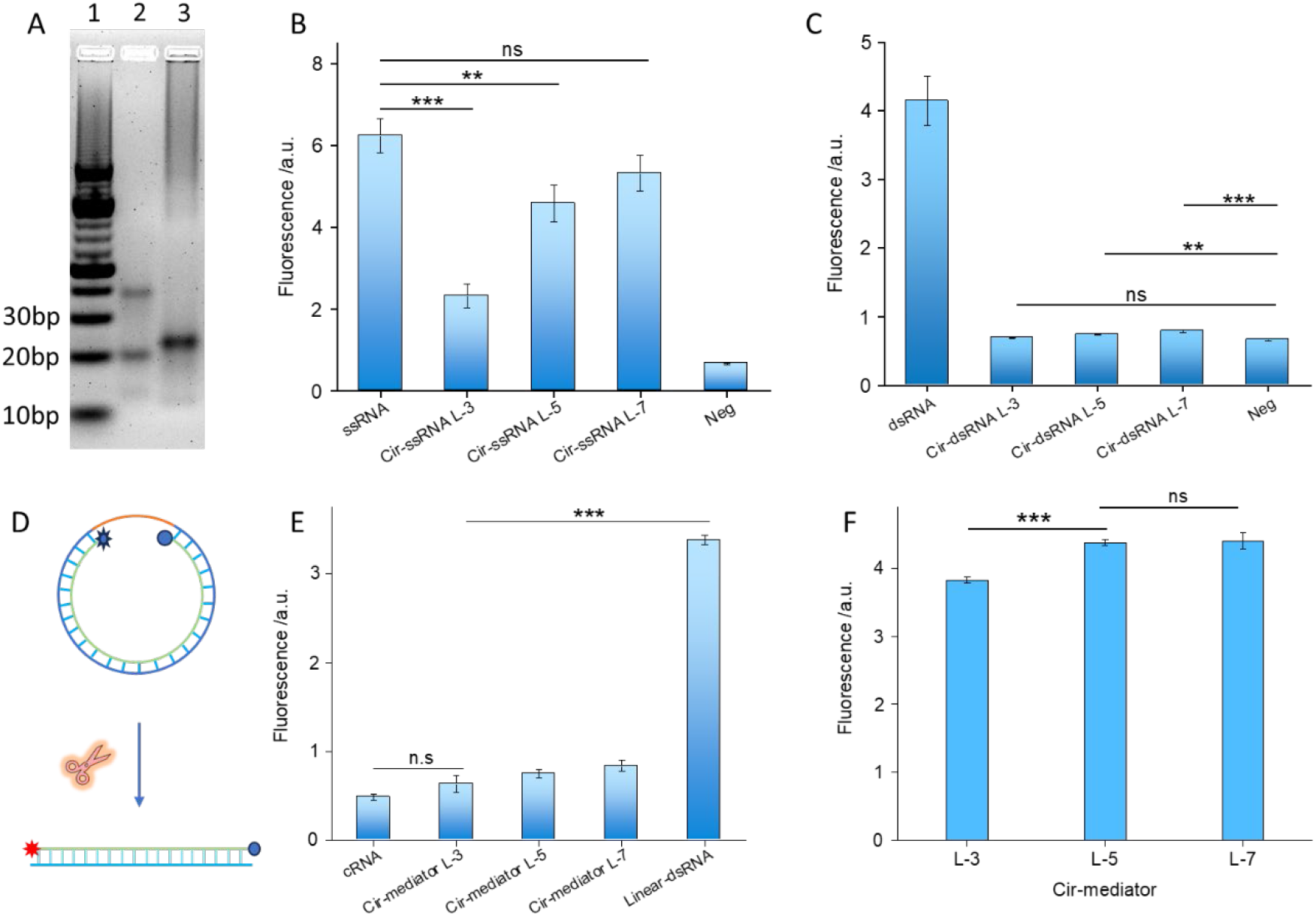
Characterization of the Cir-mediator. (A) Evidence of circular single-stranded RNA formation through gel electrophoresis assay. 1). 10 bp ladder; 2). Linear ssRNA; 3) Circular ssRNA. (B) Exploring the ability of circular ssRNA to trigger activation of Cas13a RNP (n=3), L-x represents the ssRNA linker length; (C) Exploring the ability of circular dsRNA to trigger activation of Cas13a RNP (n=3), L-x represents the ssRNA linker length; (D) Schematic of fluorescent Cir-mediator; (E) Background signal of fluorescent Cir-mediator (n=3), L-x represents the ssRNA linker length; (F) Investigation of the linker length of fluorescent Cir-mediator in a CRISPR/Cas13a biosensing system (n=3). (* P<0.05, ** P< 0.005, *** P< 0.001)

To open the locked circular targets, a fluorescent Cir-mediator was established (Fig. 3D), where a fluorescent reporter (FAM) and a quencher (BHQ1) were placed in close proximity on the Cir-mediator on both sides of the cRNA. Following the cleavage of the ssRNA region in the Cir-mediator, a linearised dsRNA is produced, which unquenches the fluorophore leading to restoration of fluorescence signals. The background fluorescence signal of different fluorescent Cir-mediators with varying linker length was tested (Fig. 5E). We found that longer ssRNA linker length leads to higher fluorescence background due to increased fluorophore-quencher distance. Subsequently, the fluorescent Cir-mediator was applied as a standard reporter in a CRISPR/Cas13a biosensing system (Fig. 5F). We found that following the addition of trigger ssRNA, the fluorescence signal of the Cir-mediator-based CRISPR/Cas13a biosensing system continued to increase with increasing incubation time, indicating the feasibility of biosensing (Fig. S5). Moreover, it was observed that increasing the linker length resulted in stronger signals, with L-5 identified as the optimal length (Fig. 5F). The stability of fluorescent Cir-mediator was also investigated (Fig. S6), and we found that the Cir-mediator shows very high stability in the rCutSmart buffer and acceptable stability in human plasma within 15min.

### 4. Assessing the fundamental characteristics of Cas13a-based autocatalytic biosensor

The operation of a Cir-mediator-based Cas13a autocatalytic biosensor is illustrated in Fig. 4A. After introducing a target RNA, Cas13a RNP is activated to *trans*-cleave the ssRNA region of the Cir-mediators. The linearized Cir-mediators become surrogate targets able to activate new Cas13a RNP and sustain an autocatalytic loop^18^. We evaluated the biosensing performance of this Cas13a-autosensor in Fig. 4B and 4C. In comparison with a basic standard Cas13a biosensor, which shows a linear signal increase with increasing time, the Cas13a-autosensor shows an exponential signal increase (Fig. 4B), and it achieves the limit of detection of 1aM (Fig. 4C).

**Figure 4.**
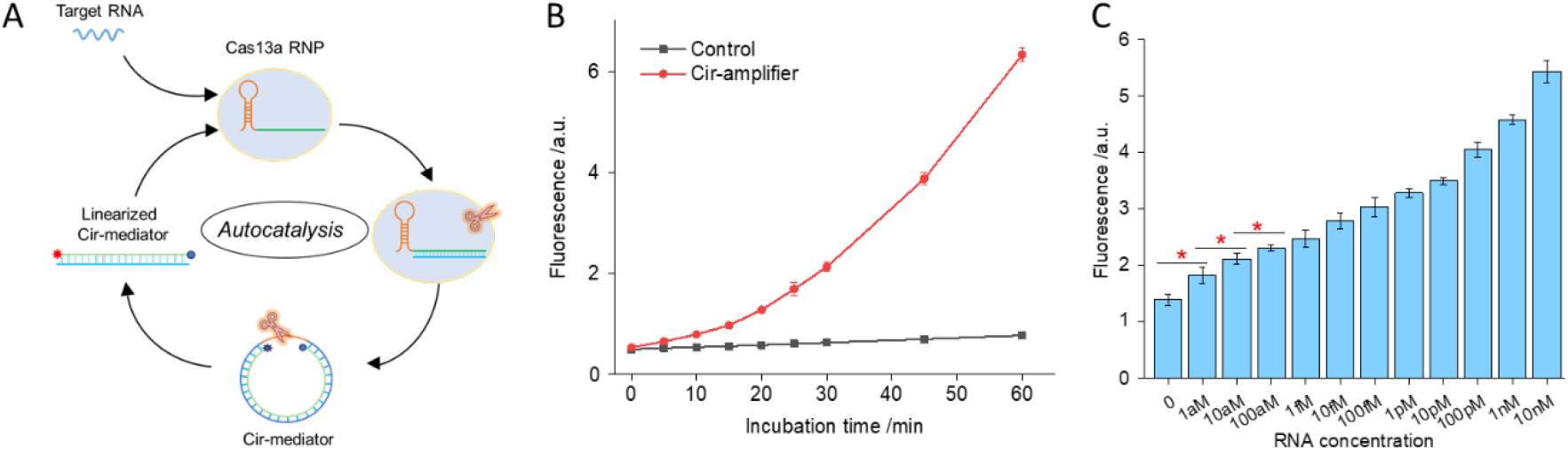
Evaluating the basic properties of Cas13a-autosensor (Cas13a-autosensor-1). (A) Mechanism of a Cas13a-autosensor; (B) Verification of autocatalysis in the Cas13a-autosensor (n=3); (C) Investigation of the sensitivity of Cas13a-autosensor using short synthetic RNA targets (n=3). (* P<0.05, ** P< 0.005, *** P< 0.001)

### 5. Application of Cas13a-based autocatalytic biosensor to clinical samples

Although single Cas RNP based Cas13a-autosensor (Cas13a-autosensor-1) realized the limit of detection of 1aM, its versatility was restricted to one specific RNA target matching the Cir-mediator sequence. To improve the versatility, we devised a two Cas RNPs based autocatalytic system (Cas13a-autosensor-2, Fig. 5A). In this system, first the target RNA activates Cas13a RNP-1 to *trans*-cleave Cir-mediators matching gRNA in RNP-2 present in its proximity. This then produces linearized Cir-mediators able to trigger activation of Cas13a RNP-2, which continue to *trans*-cleave the Cir-mediators and drive the autocatalytic reaction system. In contrast to a single Cas13a-autosensor (Fig. 4A), the two Cas13a RNP based autosensor (Fig. 5A) is able to detect different targets by only changing the gRNA of Cas13a RNP-1, but using the same Cir-mediators matching gRNA in Cas13a RNP-2.

**Figure 5.**
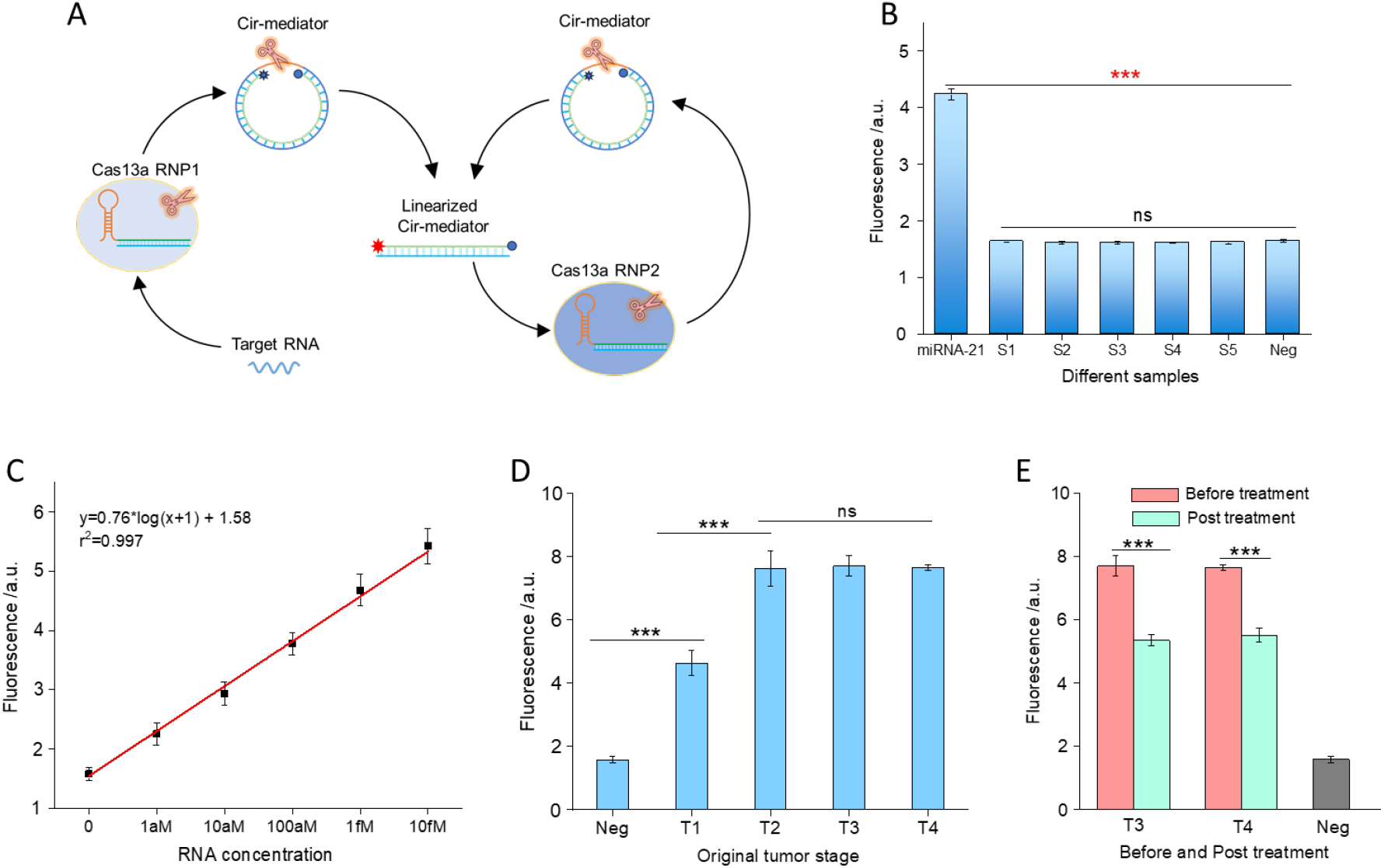
The application of Cas13a based autocatalytic biosensor (Cas13a-autosensor-2). (A) Schematics of Cir-mediator assisted autocatalytic biosensing system using two Cas13a RNPs (Cas13-autosensor-2); (B) Investigation of the specificity of Cas13a-autosensor-2 for different interference sequences (n=3); (C) The calibration curve of Cas13a-autosensor-2 for the detection of miRNA-21 in human plasma; (D) Testing the concentration of miRNA-21 in different tumor stages (Neg: 0; T1: 1.2fM; T2: 10.2pM; T3: 13.3pM; T4: 11.4pM); (E) Testing the concentration of miRNA-21 before and post treatment (T3 post treatment: 10.2 fM; T4 post treatment: 17.1 fM), Neg represents healthy human plasma. (* P<0.05, ** P< 0.005, *** P< 0.001)

To investigate whether any interfering sequences are able to activate the Cas13a-autosensor-2, a range of different sequences were added into Cas13a-autosensor-2 solution, and significant differences were observed between matched and mismatched targets, demonstrating exceptional specificity of Cas13a-autosensor-2 (Fig. 5B). Subsequently, the optimized Cas13a-autosensor-2 was applied to detect miRNA-21 in human colorectal cancer (CRC) plasma samples. Earlier reports indicate that miRNA-21 in plasma represents a potential clinical CRC biomarker which has 90% specificity and sensitivity^20^, and its physiological values in healthy and CRC patients range from fM to nM^21^. We established the calibration curve by spiking different concentrations of miRNA-21 in healthy human plasma (Fig. 5C). The Cas13a-autosensor-2 was then applied to test the miRNA-21 concentrations at different tumor stages (Fig. 5D and Table S1), and significantly higher concentration of miRNA-21 was observed in all the CRC plasma samples (T1 to T4) in comparison with healthy human plasma (Neg). In addition, we observed that the concentration of miRNA-21 rises from femtomolar (fM) to picomolar (pM) levels as the tumour stage progresses from T1 to T2. Subsequently, it stabilizes within the picomolar range from T2 to T4. Post chemotherapy treatment, the concentration of miRNA-21 was found to be reduced from pM range to fM range (Fig. 5E). Taken together, these data confirm that Cas13a-autosensor is an effective approach for the monitoring of CRC progression and treatment.

## Discussion

In this study we discovered new fundamental properties of Cas13a that dsRNA effectively triggers activation of Cas13a RNP, but activated Cas13a RNP is unable to cleave dsRNA targets. Circular RNA which has a dsRNA and ssRNA section (Cir-mediator) with optimised lengths has limited ability to activate Cas13a, but activation is restored when circular RNA is linearised by *trans*-cleavage. Based on this finding, we established a Cir-mediator-assisted Cas13a autocatalytic biosensing system (Fig. 4A). The new design changes the original activation pattern from one-to-one correspondence between RNA targets and activated Cas RNPs, to a single target activating an avalanche of activated Cas RNPs. This elegant design of a Cas13a-autosensor has been able to detect 1aM targets within 15min (Fig. 4). It provides a novel platform for ultrasensitive and rapid detection of RNA targets without preamplification.

Previous reports of ultrasensitive detection of RNA target had to introduce signal amplification techniques to integrate with CRISPR/Cas system^9-12^, or, alternatively, different Cas tandem systems were developed^13-16^. These signal amplification techniques are difficult to implement in a point-of-care setting. The tandem systems were able to realize aM level sensitivity with 2-3 hour incubation time^3, 15^ or fM level sensitivity with 60 min incubation time^14, 16^. However, for a POCT setting, aM level sensitivity with less than 15-20 min incubation time is expected. The optimized Cas13a-autosensor realized the limit of detection of 1aM within 15min, demonstrating its capability to realize PCR level sensitivity in a POCT setting.

To further improve the versatility of the Cas13a-autosensor, a two Cas13a RNP based autocatalytic sensing system was established (Fig. 5A), in which Cas13a RNP-1 recognizes the target RNA, and Cas13a RNP-2 and matching Cir-mediators are responsible for the autocatalysis reaction. In contrast to a single Cas13a-based autosensor, which is only applicable to one specific target since the dsRNA trigger in the Cir-mediator is identical to the genuine target RNA, the two Cas13a RNP-based autosensor is able to detect diverse targets by a simple replacement of gRNA of the Cas13a RNP-1. This two Cas13a RNP system transforms the standard Cas13a reaction system into an autocatalytic system with the addition of a universal Cas13a autosensor solution (Cas13a RNP2 and matching Cir-mediator). The combination of Cir-mediators and Cas13a RNP-2 mixtures can serve as an autonomous signal amplifier for all existing Cas13-based biosensing systems^22^. The optimized Cas13a-autosensor was successfully applied to monitor the miRNA-21 concentration from clinical CRC plasmas, demonstrating clinical applicability of this autosensor. The implementation of such assays in clinical practice would give the practitioners and patients much increased certainty that clinical decisions are optimally made and based on the state-of-the art clinical evidence.

## Online Methods

### 1. Materials

LwCas13a (Magigen), Cas13a reaction buffer (Magigen), rCutSmart Buffer (NEB), NEBuffer 2.1 (NEB), agarose (ThermoFisher), SYBR Green RNA dye (ThermalFisher), 10 bp DNA ladder (ThermoFisher), 6X RNA loading dye (ThermoFisher), exonuclease T (New England Biolab), copper sulfate (CuSO_4_) (Sigma, 209198), Tris(2-carboxyethyl) phosphine (TCEP) (Sigma, C4706), tris(benzyltriazolylmethyl) amine (TBTA) (ChemSupply, T2993), streptavidin coated magnetic particles (Spherotech, SVM-08-10), DNase/RNase free water (ThermoFisher), and phosphate buffered saline (PBS) (Sigma, 10 mM, pH=7.4).

All DNA and RNA oligos are synthesized and modified by Sangon Bio-Tech Ltd.

**Table 1.**
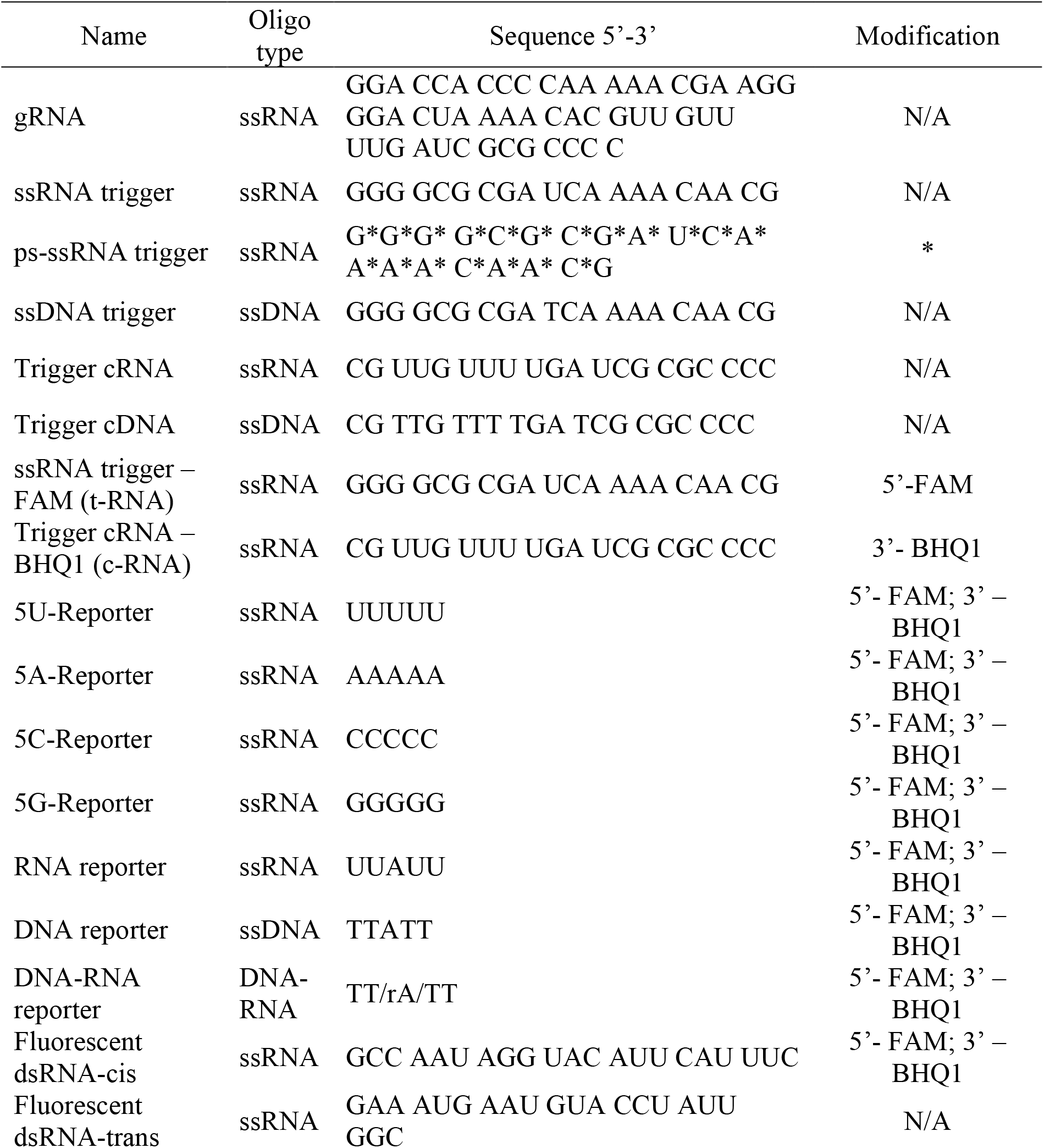
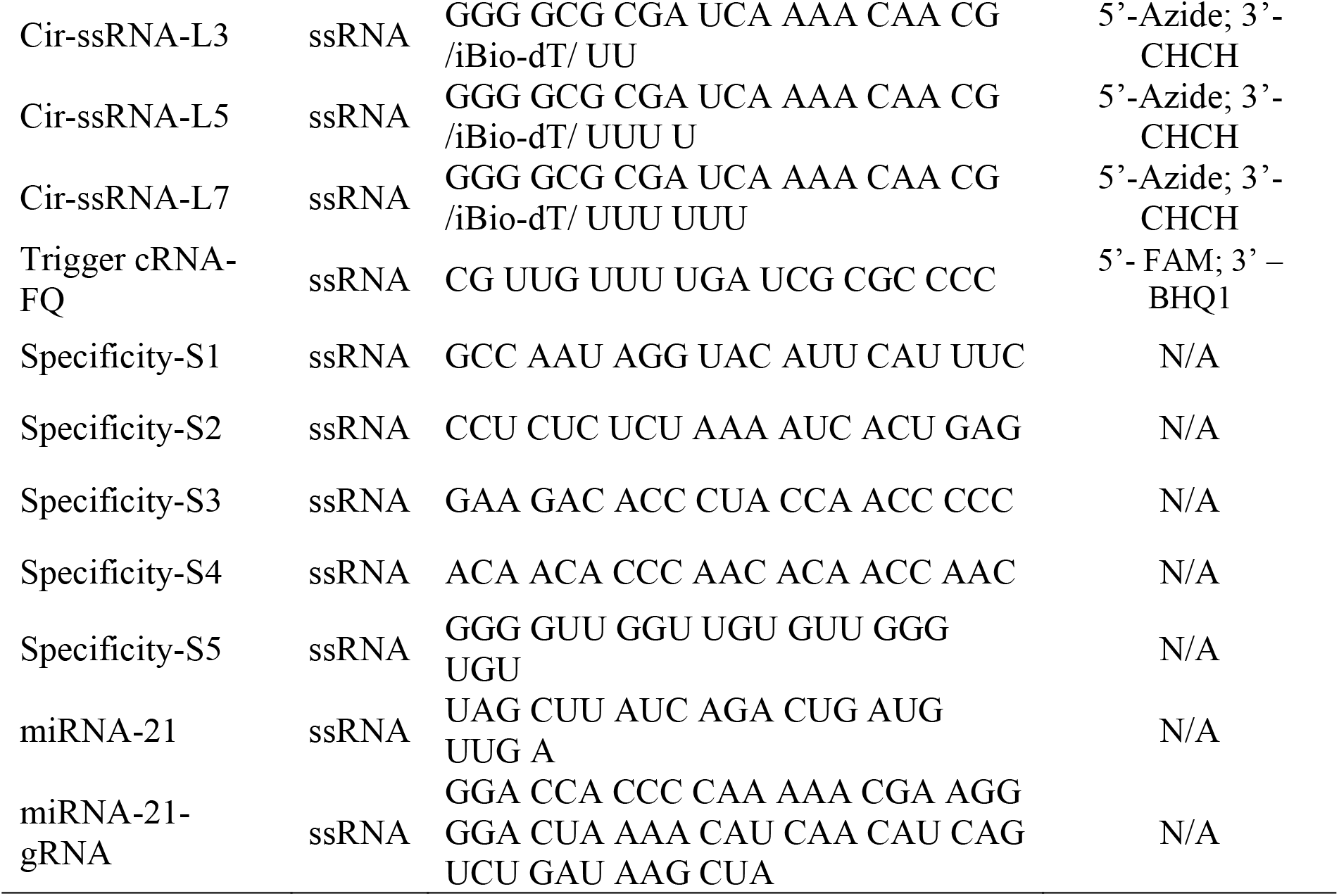
DNA and RNA oligos used in this study.

### 2. Investigation of the basic properties of single strand trigger for Cas13a RNP

Standard CRISPR/Cas13a reaction mixture was prepared by combining 40nM of Cas13a protein, 20nM of gRNA, 120nM of 5U-reporter, and 1mL of rCutSmart buffer.

Investigation of different single strand trigger types was carried out as follows. 2μL of 1μM of trigger (ssRNA, ssDNA, and ps-ssRNA) was added into 100μL standard reaction mixture and incubated at 37°C for two hours. The fluorescence signal was tested using an ID5 plate reader (Ex 480nM, and Em 520nM).

### 3. Investigation of the basic properties of double strand trigger for Cas13a RNP

Standard CRISPR/Cas13a reaction mixture was prepared by combining 40nM of Cas13a protein, 20nM of gRNA, 120nM of 5U-reporter, and 1mL of rCutSmart buffer.

Investigation of different types of double strand trigger was carried out as follows. 2μL of 1μM of trigger (ssRNA, cRNA, dsRNA, cDNA, ssRNA/cDNA, dsDNA, ps-ssRNA/cDNA) was added into 100μL of standard reaction mixture and incubated at 37°C for two hours. The fluorescence signal was tested using an ID5 plate reader (Ex 480nM, and Em 520nM).

### 4. Investigation of trigger mechanism of dsRNA trigger for Cas13a RNP

To investigate the trigger activation mechanism of Cas13a, the FRET method was utilized. In brief, a reaction mixture was prepared by combining 40nM of Cas13a protein, and 20nM of gRNA in 1mL of rCutSmart buffer. Afterwards, 2μL of 1μM of fluorescent dsRNA was added into 100μL of the prepared reaction mixture. Subsequently, the fluorescence signal was tested using an ID5 plate reader (Ex 480nM, and Em 520nM). For comparison, 2μL of 1μM of dsRNA or 2μL of 1μM of ssRNA-FAM was added into 100μL of 1X rCutSmart buffer, and the fluorescence signal was tested using an ID5 plate reader (Ex 480nM, and Em 520nM).

### 5. Investigation of the basic *trans*-cleavage properties of Cas13a RNP

To investigate the *trans*-cleavage properties of Cas13a RNP on a single strand nucleic acid, a CRISPR/Cas13a reaction mixture was first prepared comprising 40nM of Cas13a protein, 20nM of gRNA, and 120nM of reporter (RNA, DNA, RNA-DNA, 5A, 5U, 5C, and 5G) in 1mL of rCutSmart buffer. Subsequently, 2μL of 1μM of ssRNA trigger was added into 100μL standard reaction mixture and incubated at 37°C for two hours. The fluorescence signal was tested using an ID5 plate reader (Ex 480nM, and Em 520nM).

To investigate the *trans*-cleavage properties of Cas13a RNP on a double strand nucleic acid, a CRISPR/Cas13a reaction mixture was first prepared comprising 320nM of Cas13a protein, 160nM of gRNA, and 1μM of dsRNA target in 1mL rCutSmart buffer. Subsequently, 16μL of 1μM ssRNA trigger was added into 100μL standard reaction mixture and incubated at 37°C for two hours. Afterwards, agarose gel electrophoresis was applied to evaluate the conformation of dsRNA. In brief, 5% agarose gel in 1×TBE buffer was prepared with SYBR Green DNA dye. 10 μL of dsRNA was premixed with 2 μL of 6X DNA gel loading dye and then loaded into gel for electrophoresis, which was carried out for 40 min at a constant voltage of 100V. 5 μL of 10 bp DNA ladder was used for molecular weight reference. Gel images were visualized by using a Gel Doc + XR image system (Bio-Rad Laboratories Inc., USA).

To further investigate the *trans*-cleavage properties of Cas13a RNP on a fluorescent double strand nucleic acid, a CRISPR/Cas13a reaction mixture was first prepared comprising 40nM of Cas13a protein, 20nM of gRNA, and 120nM of fluorescent dsRNA reporters in 1mL of rCutSmart buffer. Subsequently, 2μL of 1μM of ssRNA trigger was added into 100μL standard reaction mixture and incubated at 37°C for two hours. The fluorescence signal was tested using an ID5 plate reader (Ex 480nM, and Em 520nM).

### 6. Synthesis and characterization of circular RNA

The synthesis approach of circular RNA was based on our previous approach (click chemistry)^18^. In brief, 0.4 mL of 0.5% w/v of streptavidin modified magnetic beads (0.74 μm) were first blocked with 1% BSA solution for 1 h to eliminate non-specific binding. Afterwards, 1 mL of 0.5 μM biotinylated linear-ssRNA was incubated with the beads for 1 h following a PBS wash to remove the residual free linear-ssRNA. Subsequently, 1 mL of the click chemistry reaction solution (1.0 mM of CuSO_4_, 2.0 mM of TCEP, and 100 μM of TBTA) was added and incubated with the beads for 12 h at room temperature. After synthesis, the magnetic beads were collected and washed with PBS buffer to remove excess chemicals. Subsequently, 100 μL of 100 units/mL of Exonuclease T solution was added and incubated at 37 °C for 30 min to remove linear ssDNA. After washing with PBS buffer, the synthesized Cir-ssRNA was released from the streptavidin-modified magnetic beads by heat treatment at 95°C for 30 min, and the supernatant was collected for further use. All the Cir-ssRNA used in this research are synthesized based on this approach. A Nanodrop system was utilized to test the concentration of synthesized Cir-ssDNA.

The formation of Cir-ssRNA was verified by using agarose gel electrophoresis. In brief, 5% agarose gel in 1×TBE buffer was prepared with SYBR Green DNA dye. 10 μL of Cir-ssRNA was premixed with 2 μL of 6X DNA gel loading dye and then loaded into gel for electrophoresis, which was carried out for 40 min at a constant voltage of 100V. 5 μL of 10 bp DNA ladder was used for molecular weight reference. Gel images were visualized by using Gel Doc + XR image system (Bio-Rad Laboratories Inc., USA).

### 7. Investigation of the basic trigger ability of circular RNA to Cas13a RNP

To investigate the trigger ability of Cir-ssRNA and Cir-dsRNA, standard CRISPR/Cas13a reaction solution was first prepared (40nM of Cas13a protein, 20nM of gRNA, 120nM of reporter, and 1mL of rCutSmart buffer). Subsequently, 2μL of 1μM Cir-ssRNA or Cir-dsRNA trigger was added into 100μL of the standard reaction mixture and incubated at 37°C for two hours. The fluorescence signal was tested using an ID5 plate reader (Ex 480nM, and Em 520nM).

### 8. Investigation of the *trans*-cleavage properties of Cas13a RNP on Cir-mediator

The formation of fluorescent Cir-mediator was conducted by mixing synthesized Cir-ssRNA with its corresponding fluorescent cRNA at the molar ratio of 1:1. Afterwards, the background of fluorescent Cir-mediator (120nM) in rCurSmart buffer was tested using an ID5 plate reader (Ex 480nM, and Em 520nM).

To investigate the *trans*-cleavage properties of Cas13a RNP on Cir-mediator, a CRISPR/Cas13a reaction mixture was first prepared comprising 40nM of Cas13a protein, 20nM of gRNA, and 120nM of fluorescent Cir-mediator in 1mL of rCutSmart buffer. Subsequently, 2μL of 1μM ssRNA trigger was added into 100μL standard reaction mixture and incubated at 37°C for two hours. The fluorescence signal was tested using an ID5 plate reader (Ex 480nM, and Em 520nM).

To investigate the stability of Cir-mediator, 200 nM of fluorescent Cir-mediator (L-5) was incubated in rCurSmart buffer, 10% human serum solution, or 10% saliva solution for two hours. The fluorescence signal was tested using an ID5 plate reader (Ex 480nM, and Em 520nM).

### 9. Evaluating the basic properties of Cas13a based autocatalytic biosensor

Cas13a-autosensor-1 standard reaction mixture was first prepared: 40nM Cas13a protein, 20nM gRNA, and 120nM of fluorescent Cir-mediator in 1mL rCutSmart buffer. Subsequently, 2μL of 1μM ssRNA trigger was added into 100μL standard reaction mixture and incubated at 37°C for two hours. The fluorescence signal was tested using ID5 plate reader (Ex 480nM, and Em 520nM).

To investigate the autocatalysis activity, 1 pM ssRNA trigger was added into 100μL standard reaction mixture and incubated at 37°C for one hour. The fluorescence signal was tested using ID5 plate reader (Ex 480nM, and Em 520nM).

To investigate the sensitivity, different concentrations of ssRNA (0-10nM) was added into 100μL of the standard reaction mixture and incubated at 37°C for two hours. The fluorescence signal was tested using an ID5 plate reader (Ex 480nM, and Em 520nM).

### 10. Application of Cas13a-based autocatalytic biosensor for the detection of miRNA-21 in healthy human plasma and clinical human plasma samples

The Cas13a-autosensor-2 reaction mixture for miRNA-21 detection was prepared as follows: 40nM of Cas13a protein was mixed with 20nM gRNA (miRNA-21) to form the Cas13a RNP1. In the meantime, 40nM of Cas13a protein was mixed with 20nM of gRNA (Cir-mediator) to form the Cas13a RNP2. Afterwards, the prepared Cas13a RNP1 and Cas13a RNP2 were mixed with 120nM of fluorescent Cir-mediator in 1mL of rCutSmart buffer to form the reaction mixture. The reaction mixture was stored at 4 °C before use.

To investigate the specificity, 2μL of 1μM of interference ssRNA was added into 100μL reaction mixture for activating *trans*-cleavage of Cas13a and enabling the CRISPR/Cas biosensing reaction (37°C). The fluorescence signal was tested using an ID5 plate reader (Ex 480nM, and Em 520nM).

To establish the calibration curve, 10μL of spiked in sample was added into 90μL standard reaction mixture for activating *trans*-cleavage of Cas13a and enabling the CRISPR/Cas biosensing reaction (37°C). The fluorescence signal was tested using an ID5 plate reader (Ex 480nM, and Em 520nM).

To evaluate the clinical application performance of Cas13a-autosensor, colorectal cancer (CRC) clinical samples were collected and provided by Health Precincts Biobank of UNSW. All patients provided written informed consent for participation in this study. These plasma samples were collected from patients with different tumor stages from T1 to T4. In addition, post chemotherapy (post sx) samples from T2 to T4 were also collected to evaluate the effectiveness of treatment. For the testing, 10μL of CRC plasma samples was added into 90μL standard reaction mixture for activating *trans*-cleavage of Cas13a and enabling the CRISPR/Cas biosensing reaction (37°C). The fluorescence signal was tested using ID5 plate reader (Ex 480nM, and Em 520nM).

All human plasma experiments were approved by the UNSW Ethics Committee (UNSW HC210160). The Health Precincts Biobank samples were received from operates under ethics approval from the South Eastern Sydney Local Health District – Northern Sector HREC, reference number 2019/ETH12304.

## Notes

### Competing Interest Statement

The authors have declared no competing interest.

