## Supplemental file for "Modulating Cas13a *trans*-cleavage by double strand RNA: Application to the development of an autocatalytic sensor"

### Supplementary methods

#### 1. Investigation of the basic properties of CRISP/Cas13a biosensing system

Standard CRISPR/Cas13a reaction mixture consisted of 40nM of Cas13a protein, 20nM of gRNA, 120nM of 5U-reporters, and 1mL of rCutSmart buffer.

Optimization of trigger ssRNA concentration: A variety of trigger ssRNA (0, 2.5, 5, 10, and 20 nM) were added into 100µL of a standard reaction mixture and incubated at room temperature for two hours. The fluorescence signal was tested using an ID5 plate reader (Ex 480nM, and Em 520nM).

Optimization of gRNA to Cas13a ratio: A variety of gRNA-based reaction mixture was prepared (0, 5, 10, 20, 40, and 80nM of gRNA), 40nM of Cas13a protein, 120nM of reporters, and 1X of rCutSmart buffer. Afterwards, 40nM of trigger ssRNA was added into 100µL of a standard reaction mixture and incubated at room temperature for two hours. The fluorescence signal was tested using an ID5 plate reader (Ex 480nM, and Em 520nM).

Optimization of reporter concentration: A variety of 5U-reporter solutions was prepared (0, 20, 40, 60, 120, 180 nM of reporter), 40nM of Cas13a protein, 20nM of gRNA, and 1X of rCutSmart buffer. Afterwards, 20 nM of trigger ssRNA was added into 100µL of the standard reaction mixture and incubated at room temperature for two hours. The fluorescence signal was tested using an ID5 plate reader (Ex 480nM, and Em 520nM).

Optimization of buffers: Three different types of buffers were used as the reaction buffer, including Reaction buffer, NEB2.1, and rCutSmart buffer. After preparation of the standard reaction mixture, 20 nM of trigger ssRNA was added into 100µL of the standard reaction mixture and incubated at room temperature for two hours. The fluorescence signal was tested using an ID5 plate reader (Ex 480nM, and Em 520nM).

Optimization of temperature: After preparation of the standard reaction mixture, 20 nM of trigger ssRNA was added into 100µL of the standard reaction mixture and incubated at room temperature or 37°C for two hours. The fluorescence signal was tested using anID5 plate reader (Ex 480nM, and Em 520nM).

Investigation of the limit of detection of standard CRISPR/Cas13a biosensing system: After preparation of the standard reaction mixture, a variety of trigger ssRNA solutions (0, 1pM, 10pM, 100pM, 1nM, 10nM, and 100nM) were added into 100µL of the standard reaction mixture and incubated at room temperature for two hours. The fluorescence signal was tested using an ID5 plate reader (Ex 480nM, and Em 520nM).

### Supplementary figures

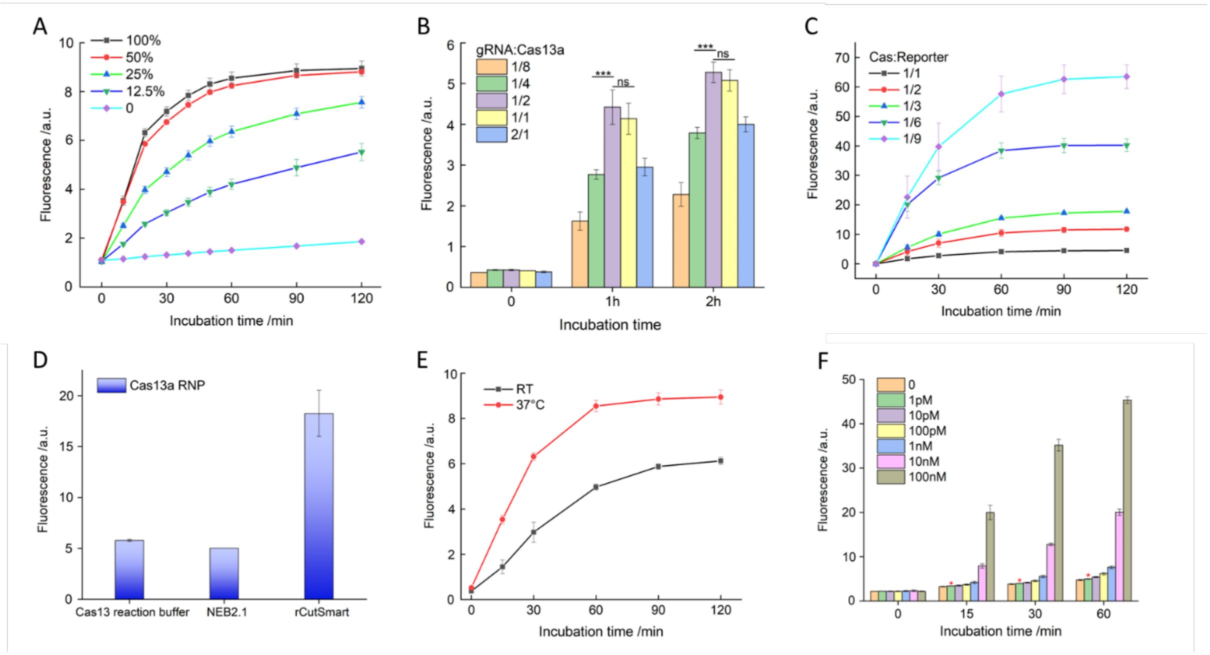


**Figure S1**. Investigation of the basic properties of CRISPR/Cas13a biosensing system. (A) Investigation of ssRNA trigger to Cas13a ratio. We found that 100% trigger provides the highest fluorescence signal; (B) Investigation of gRNA to Cas13a ratio. We found the optimum gRNA to Cas13a ratio is 1:2; (C) Investigation of reporter ratio, and higher reporter concentration leading to better performance; (D) Investigation of different types of reaction buffers. We found that rCutSmart shows optimum performance; (E) Investigation of reaction temperature. 37 °C was found to be the optimal.; (F) Investigation of the biosensing performance of optimized CRISPR/Cas13a biosensing system. the limit of detection of 1 pM was achieved in 15min. (* $P<0.05$, ** $P< 0.005$, *** $P< 0.001$)

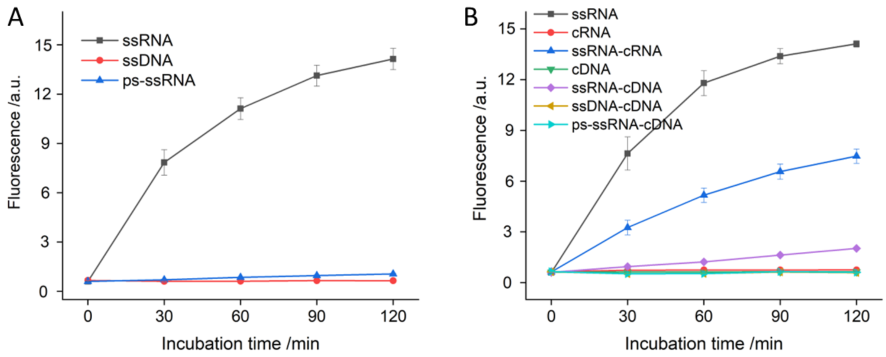


**Figure S2**. Investigation of different types of triggers for Cas13a RNP as listed in the figure. (A) Investigation of different types of single strand trigger; (B) Investigation of different types of double strand trigger

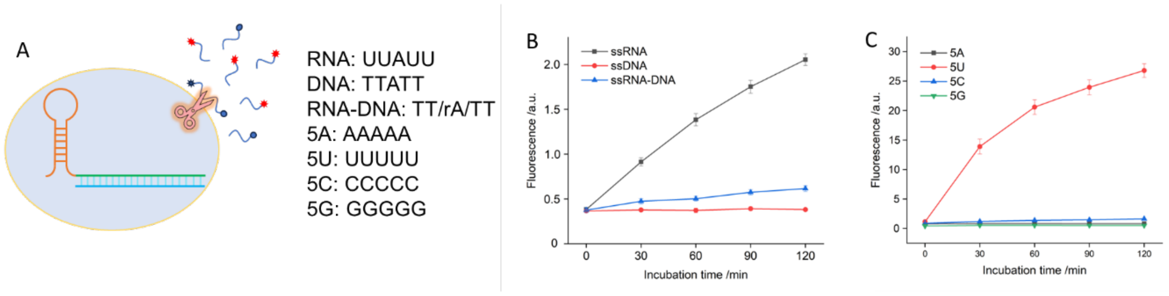


**Figure S3**. Investigation of the *trans*-cleavage substrates of Cas13a RNP as listed in the figure. (A) Schematic of the *trans*-cleavage activity of Cas13a RNP; (B) Different types of single strand substrates; (C) Different nucleobase based ssRNA substrates. (* P<0.05, ** P< 0.005, *** P< 0.001)

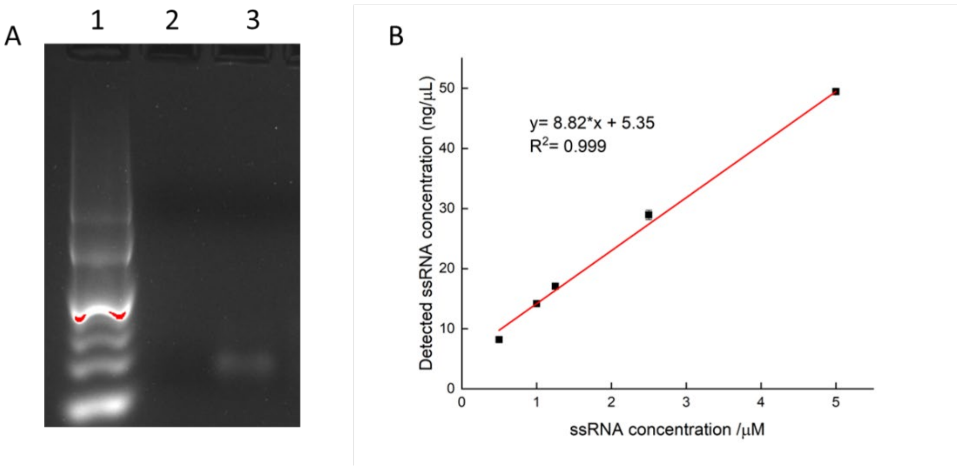


**Figure S4**. Characterization of the Cir-mediator. (A) Demonstrating the function of Exo-T on linear ssRNA. 1). 10 bp ladder; 2). Linear ssRNA with Exo T; 3) Linear ssRNA. Exo T is able to cleave linear ssRNA. (B) Calibration curve for the synthesized circular ssRNA.

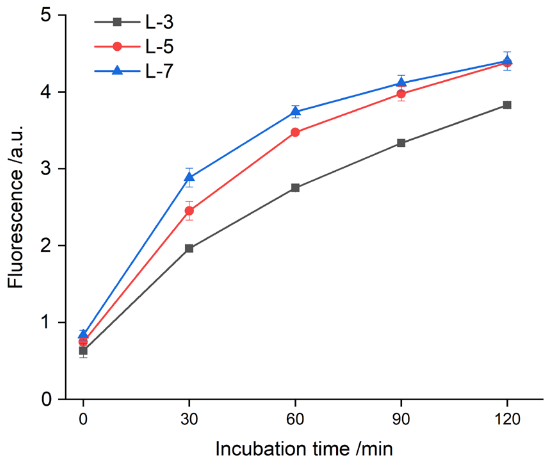


**Figure S5**. Investigation of the linker length of Cir-mediator in a CRISPR/Cas13a biosensing system (n=3).

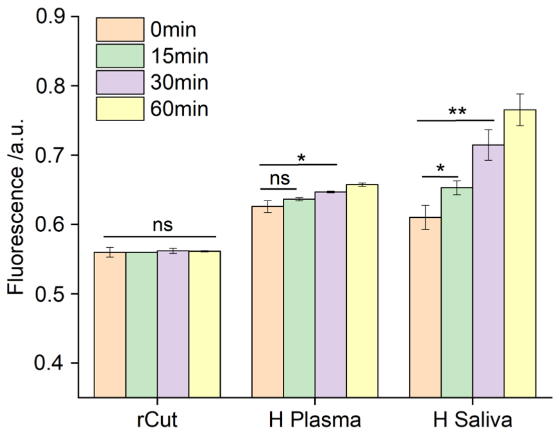


**Figure S6**. Investigation of the stability of Cir-amplifier in rCutSmart buffer, human plasma, and human saliva (n=3). (* P<0.05, ** P< 0.005, *** P< 0.001)

### Supplementary tables

**Table S1.** Clinical information of Health Precincts Biobank samples

| Participant | Consent | Specimen type | Diagnosis | Original tumor stage | Chemotherapy |
|---|---|---|---|---|---|
| HSA3151 | CONSENTED | Plasma | C20 Malignant neoplasm of rectum | T1 | No |
| HSA2313 | CONSENTED | Plasma | C19 Malignant neoplasm of rectosigmoid junction | T2 | No |

| | | | | | |
|---|---|---|---|---|---|
| HSA5907 | CONSENTED | Plasma | C18.2 Malignant neoplasm of ascending colon | T2 | No |
| HSA3135 | CONSENTED | Plasma | C20 Malignant neoplasm of rectum | T3 | No |
| HSA2459 | CONSENTED | Plasma | C20 Malignant neoplasm of rectum | T3 | No |
| HSA2699 | CONSENTED | Plasma | C18.4 Malignant neoplasm of transverse colon | T4 | No |
| HSA4423 | CONSENTED | Plasma | C19.9 Malignant neoplasm of rectosigmoid junction | T3 | Post sx |
| HSA2548 | CONSENTED | Plasma | C20 Malignant neoplasm of rectum | T3 | Post sx |
| HSA2336 | CONSENTED | Plasma | C20 Malignant neoplasm of rectum | T3 | Post sx |
| HSA4444 | CONSENTED | Plasma | C20 Malignant neoplasm of rectum | T3 | Post sx |
| HSA3067 | CONSENTED | Plasma | C18.8 Overlapping malignant lesion of colon | T4 | Post sx |
